# Noise-Cancelling Repeat Finder: Uncovering tandem repeats in error-prone long-read sequencing data

**DOI:** 10.1101/475194

**Authors:** Robert S. Harris, Monika Cechova, Kateryna D. Makova

## Abstract

**Summary:** Tandem DNA repeats can be sequenced with long-read technologies, but cannot be accurately deciphered due to the lack of computational tools taking high error rates of these technologies into account. Here we introduce Noise-Cancelling Repeat Finder (NCRF) to uncover putative tandem repeats of specified motifs in noisy long reads produced by Pacific Biosciences and Oxford Nanopore sequencers. Using simulations, we validated the use of NCRF to locate tandem repeats with motifs of various lengths and demonstrated its superior performance as compared to two alternative tools. Using real human whole-genome sequencing data, NCRF identified long arrays of the (AATGG)_n_ repeat involved in heat shock stress response.

**Availability and implementation:** NCRF is implemented in C, supported by several python scripts. Source code, under the MIT open source license, and simulation data are available at https://github.com/makovalab-psu/NoiseCancellingRepeatFinder, and also in bioconda.

## 1 Introduction

Long tandem repeat (TR) arrays are associated with heterochromatin and play critical roles in the human genome. For instance, (TTAGGG)_n_repeats protect telomeres (Blackburn and Gall, 1978), (AATGG)_n_repeats are implicated in heat shock response (Goenka *et al.*, 2016), and alpha satellites with 171-mer repeated units bind histones participating in chromosome segregation during cell division (Masumoto *et al.*, 2004). The lengths of heterochromatin-associated TRs differ across populations (Altemose *et al.*, 2014;Wevrick and Willard, 1989) and change with aging and environmental exposure (Zhang *et al.*, 2015; Goenka *et al.*, 2016). Despite these important features of TRs, their length variation has been understudied due to the lack of experimental and computational techniques that can capture their full length.

Long TRs cannot be studied with short sequencing reads, but can be profiled with long-read technologies (Pacific Biosciences, or PacBio, and Oxford Nanopore, or Nanopore). However they are difficult to decipher because such technologies have high error rates and error profiles different from those of short-read Illumina sequencing (Schirmer *et al.*, 2016). To our knowledge, no tool currently exists to identify TR arrays in long, error-prone sequencing reads. Tools solving similar problems, primarily developed to work with short reads or assembled genomes, have limitations when applied to this use case (Lower *et al.*, 2018). Some of them fail to model asymmetric rates of insertions vs. deletions (e.g. Tandem Repeats Finder, or TRF (Benson, 1999), and Tantan (Frith, 2011)), while others do not permit high sequencing error rates (e.g. short read mappers bwa (Li and Durbin, 2009) and bowtie (Langmead, 2010)). HMMER (Eddy, 1998) reports single instances, but not repeats, of a motif. PacmonSTR (Ummat and Bashir, 2014) requires annotations of TRs in an assembled genome, while many long TRs are present in unmapped reads (Altemose *et al.*, 2014; Novák *et al.*, 2017, 2010). General purpose aligners, e.g., Minimap2 (Li, 2018), permit parameterizations for different long-read sequencing technologies, but are not designed to find TRs.

To address the decoding of TR arrays in error-prone long sequencing reads, we developed Noise-Cancelling Repeat Finder (NCRF), which identifies arrays of user-specified TR motifs in sequencing reads or assembled genomes. NCRF supports high and asymmetric rates of short insertions and deletions observed in long-read sequencing data. As a result, its performance is superior to alternative tools.

## 2 Developing NCRF

NCRF utilizes a modified Smith-Waterman algorithm (Smith and Waterman, 1981) aligning a read to any number of tandem copies of a specified motif sequence (Fig. S1, Supplementary Note S1A). A modified version of Dynamic Programming (DP) uses only one copy of the motif but allows loops from the end back to the beginning, so that one column of the DP matrix represents any number of tandem copies of the motif. A modified scoring recurrence supports asymmetry in insertion vs. deletion rates and high prevalence of short indels (). Resulting alignments are local in the read but global in the looped motif, and can begin and end at any position in the motif. Internal post-alignment processing removes segments with high error density (Supplementary Note S1B).

The reported alignments identify sequence segments that putatively align to perfectly repeated copies of a motif (Supplementary Note S1A). However, segments containing a mix of variants of the motif, or of a similar but different motif, may also be reported. Such mixes are consistent with the known evolutionary signatures of heterochromatic repeats (Plohl et al., 2008). One can run an optional consensus filtering step to identify TR arrays with a single dominant motif, i.e. a motif whose each position matches over the array at least half the time (Supplementary Note S1C). Intervals reported for more than one motif can be identified with an optional overlap-detection step (Supplementary Note S1E).

## 3 Error profiles of long-read sequencing technologies

To guide our simulations and tune NCRF scoring parameters, we derived error rates for two common long-read sequencing technologies — PacBio and Nanopore (Supplementary Note S2). We observed whole-genome error rates of 15% with insertions being more common than deletions for PacBio (mismatches=1.7%, insertions=8.9%, deletions=4.3%), and error rates of 16% but favoring deletions for Nanopore (mismatches=4.6%, insertions=3.8%, deletions=7.7%; Table S1).

## 4 Analysis of simulated reads and tool comparison

We next simulated PacBio and Nanopore sequencing reads for a mock genome containing specific TRs with periods 5, 10, 20, 40, 80, and 171, length at least 500 bp, and mean length 1,000 bp (Supplementary Note S3). Prior to filtering (Supplementary Note S1C), NCRF discovered 99% and 95% of the specified TRs in PacBio and Nanopore reads, respectively (Fig. 1A and Table S2). However, false discovery rates were frequently high, particularly for the longer motifs, justifying the need for consensus filtering. Filtering reduced false discovery rates to approximately 1% for the 5-, 10-, and 20-mers, and below 4% for longer motifs. The reduction in false discovery rates due to filtering was accompanied by a reduction in true positive rates; by about 1% for the 5-, 10-, 20-, and 40-mers, and by a higher percentage for the 80- and 171-mer.

**Figure 1.**
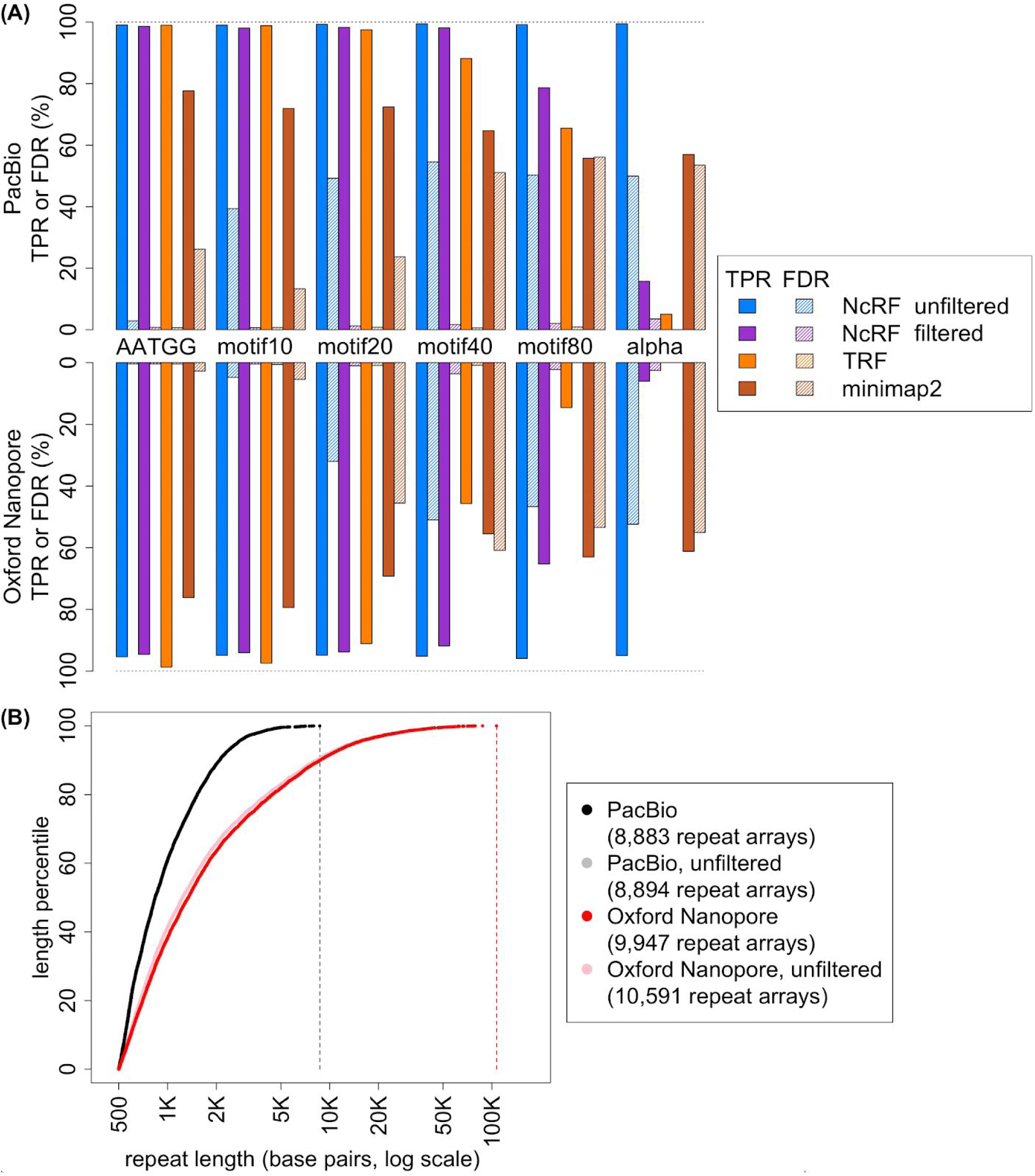
(**A**) Comparison of discovery performance of NCRF, TRF, and Minimap2 on simulated PacBio (upper panel) and Nanopore (lower panel) reads. Solid bars are true positive rates (TPR), crosshatched bars are false discovery rates (FDR). (**B**) Distribution of observed lengths of (AATGG)_n_ arrays (with and without consensus filtering) observed in PacBio and Nanopore reads. Reads were subsampled to a similar length distribution of 16.5 Gb for both technologies (see Supplementary Note S6). Filtered and unfiltered results for PacBio are very similar.

We next compared the performance of NCRF (applying consensus filtering) to that of two alternative tools — TRF (Benson, 1999) and Minimap2 (Li, 2018) (Fig. 1A, Supplementary Notes S4-S5). True positive rates were higher for NCRF than TRF for longer motifs, but were similar between the two tools for shorter motifs. TRF had low false discovery rates (less than 1%), however it failed to find any 171-mer repeats in Nanopore reads. Minimap2 had lower true positive rates and higher false discovery rates than NCRF across the board, except for true positive rates for the 171-mer. Surprisingly, both TRF and Minimap2 occasionally reported overlapping intervals for the same motif (Table S2). Several other tools were considered for this evaluation, but rejected after preliminary investigation (Supplementary Note S7).

## 5 Applying NCRF to Real Sequencing Data

Lastly, we applied NCRF to investigate perfect repeats of the (AATGG)_n_repeat in publicly available PacBio and Nanopore sequenced data (Zook *et al.*, 2016;Jain *et al.*, 2018) subsampled to a common read length distribution (Supplementary Note S6) and generated for the same individual. The resulting densities of (AATGG)_n_ for the two technologies are reported in Fig. 1B. Searching for (AATGG)_n_ repeats with length of at least 500 bp in 16.5 Gb of PacBio sequenced reads, NCRF identified 8,883 repeats. 9,766,859 bp of the reads were part of these repeats, leading to the density of 0.6 bases per kilobase sequenced. In Nanopore reads, we found 9,947 repeats, totalling 35,561 kb or 2.2 bases per kilobase. Additional applications of NCRF to real sequencing data are presented in (Cechova *et al.*, 2018), where potential reasons behind differences in density between technologies are discussed.

## 6 Conclusions

To our knowledge, NCRF is the first tool designed specifically to identify TR arrays in noisy sequencing data, accounting for the unique characteristics of the long-read technologies. We anticipate it will accelerate research of heterochromatin-associated TR arrays and will aid in unraveling their functions in the genome.

## Supporting information

## ACKNOWLEDGEMENTS

We thank Paul Medvedev and Marzia Cremona for critical reading of the manuscript.

