## Supplementary material for "Noise-Cancelling Repeat Finder: Uncovering tandem repeats in error-prone long-read sequencing data"

### SUPPLEMENTARY INFORMATION

#### SUPPLEMENTARY NOTES

##### Supplementary Note S1. NCRF Implementation

The NCRF workflow is shown in Figure S1. A list of repeat motifs and a set of sequences (reads) are input, and the output is a list of read intervals that show evidence for the motifs. End-user output is typically in tabular text, suitable as input for other tools.

The first steps in the pipeline are implemented in the C program NCRF. This performs alignment between the reads and the motifs, removes super-noisy alignment segments, and reports alignments that meet user-defined criteria for "long enough" and "not too error-laden".

Output from the C program is then post-processed by a series of python scripts. In the figure, alignments that fail consensus filtering (defined below) are removed, and the surviving alignments are reduced to a summary format.

**1A. Looping Aligner**

The aligner at the core of NCRF finds alignments of a given motif to a segment of a given DNA sequence, with the motif repeated as often as needed. It is a Smith-Waterman aligner (Smith and Waterman, 1981) with affine gap penalties. It makes use of a typical Dynamic Programming (DP) matrix with a row for each nucleotide in *a single copy* of the motif and a column for each nucleotide in the sequence. Typically, the alignment core utilizes a score for matches and penalties for mismatches and indels; but different penalties are allowed for insertions and deletions since sequencing technologies can be biased as to which type of indel they introduce.

However, the basic recurrence relation is modified in a few ways. First, to accommodate the repeated motif, the match/substitution step allows loops from the end of the motif to the beginning, i.e. from the end of one column of the DP matrix to the start of the next. Similarly, the deletion step also allows loops, from the end of a column back to its start. This requires a partial, adaptive, second pass of each column to allow a series of deletions containing such a loopback. This second pass terminates when it reaches a cell not improved by deletion; in practice the second pass terminates within the first few cells. Because we allow loops, an optimal path in the DP matrix will typically look something like a sawtooth wave, increasing the row (motif position) as it moves along the sequence, then jumping back to the first row. Each saw 'tooth' represents another copy of the motif.

The second change to the recurrence is that we disallow back-to-back insertion openings, even if the penalty to extend an insertion is worse than the penalty to open a new insertion. This reflects the nature of the targeted sequencing technologies (such as PacBio), in which single-base insertions are observed more frequently than longer insertions. Deletions are treated similarly.

Third, we allow alignments to begin and end in any position of the motif. E.g., for AATGG we may find an alignment of GGAATGGAATGGAATGGA.

And lastly, where Smith-Waterman finds only a single highest-scoring local alignment, we remove the aligning segment and repeat the process on the remainder of the sequence.

**1B. Bridges and Tails**

An often ignored side effect of the alignment optimization model is that incorporating false positive segments into an alignment can increase its score. Consider two nearby alignments having high positive scores *X* and *Y*. The intervening interval may be very noisy and thus has a negative score -*Z*. In isolation, this interval would be discarded. But if *X*>*Z* and *Y*>*Z*, the combined alignment will have score *X*+*Y*-*Z*, larger than either *X* or *Y*, and can be reported as one alignment. We call such an intervening interval an *unrelated bridge*. Note that any single error will also fit that definition; in reality we are interested in sufficiently long intervals exhibiting this property.

Further, any pair of random, unrelated DNA sequences has a non-zero probability of containing a positive-scoring prefix or suffix. This is especially true if the scoring scheme is designed to tolerate a lot of errors. Thus any correct alignment A might be extended by a false positive *unrelated tail* B, since B increases the overall score.

In NCRF, we address this problem by internally post-processing alignments to excise segments containing clumps of errors. We justify this step by our expectation of error rates to be higher in misaligned segments than in true homology.

**1C. Consensus Filtering**

NCRF aims to find near-perfect alignments of a given repeat motif. Occasionally, such repeat motifs might be interleaved with other (often similar) repeat motifs. Thus, the resulting long array will be a mix of multiple motifs. These additional motifs are expected to inflate the observed error rates.

If an alignment is truly to a near-perfect alignment of the sought motif, we should expect, at each position along the motif, to see a match to the nucleotide at that position more often than any other alignment event (i.e. a mismatch or indel). We implemented a filtering process based on that observation. Given an alignment, we collect and count what was aligned at each column. We form a consensus motif from the majority at each column, and if this consensus matches the sought motif, the alignment passes the filter. Otherwise, the consensus differs from the sought motif (or no consensus was formed because not all columns had a majority), and the alignment is discarded.

In detail, the filter slices an alignment into pieces, each piece aligning to a single perfect copy of the sought motif. From this, it effectively produces a multiple-sequence alignment containing one column for each position in the motif. Each column is reduced to its majority (if it has one), and the majorities are concatenated into a consensus motif.

The corresponding code is implemented in ncrf_concensus_filter.py.

**1D. Scoring Parameters**

The error profile for a given sequencing technology (Table S1) was applied to a simulated molecule containing 1,000 perfect repeats of AATGG with lengths sampled from a normal distribution with

$\mu = 2,500$ and $\sigma = 1,667$ (lengths below 500 bp were discarded). Repeats were separated, on average, by 500 bp of random DNA sequence. The simulator (simulate_reads.sh) produced a single read corresponding to the entirety of the molecule, with errors sampled from the given profile. It also reported the ground truth — the positions of the repeated motifs.

Knowing the ground truth in the simulated read allows us to treat the aligner, with a scoring set, as a classifier. A scoring set can then be evaluated by standard classifier metrics derived from True/False Positives/Negatives. As discussed below, early experimentation revealed that false discovery rate (FDR) varied little between scoring sets, and we concentrated on maximizing true positive rate (TPR).

The optimization process was equivalent to a short random walk with an optimized random step. At each "step", many random scoring sets in the neighborhood of the current optimum were evaluated (using parameter_ball.py). The match score was held at 100 while the five penalties were each allowed to vary in the range of +/-150 from the previous optimum. Scoring sets with any penalty below 10 were excluded. Scoring sets that would cause the aligner to prefer a back-to-back insertion/deletion pair to a mismatch were also excluded. Each set was given to NCRF to align for the motif, and the output was compared to the known truth (these steps are performed by evaluate_scoring_sets.sh). Classier performance for all sets were plotted as TPR vs. FDR, and the new optimum was selected manually. This process was repeated for only five steps, at which point the new optimum was not an improvement. This optimum was then tested for overfitting to the specific simulated read. 100 other simulated reads were generated, the scoring set was evaluated as a classifier on these reads, and the results were manually compared. The optimum resulting from this process was then installed in NCRF as the default scoring set for the corresponding sequencing technology.

The TPR achieved in this process was about 95% for both PacBio and Nanopore, using error models derived from the published alignments for HG002 (Supplementary Note S2). FDR was typically less than 0.05%. This is expected because in this simulation the true negatives consisted solely of random sequences.

The optimum scoring sets derived by this process are shown in Table S3.

**1E. Overlapping Alignments**

When only one motif is searched for, the alignments initially reported by NCRF will not overlap. However, when two or more similar motifs are searched for, some intervals of a read may align to more than one repeat. Such a situation might inflate the overall repeat density.

To address this situation, the NCRF package includes a post-processing script, ncrf_resolve_overlaps.py, that groups alignments by those unique to each motif, and those that align to some larger subset of the motifs.

#### Supplementary Note S2. Error Models

Error models for both sequencing technologies were derived from public alignments of human sequencing reads (of the same human individual) to a reference genome. For PacBio, we used alignments from (Zook *et al.*, 2016). The library for HG002 was prepared using Illumina TruSeq PCR-Free LT Sample Prep Kit, sequenced using Illumina HiSeq 2500 and subsequently aligned to GR37 using blasr (Nowak *et al.*, 2018). For Nanopore, we used alignments from (Zook *et al.*, 2016) computed by Minimap2 (Li, 2018). That project prepared HG002 DNA using modified versions of Josh Quick's protocol (Quick, 2018), and a mix of SQK-RAD003 and SQK-RAD004 library prep kits, sequenced on MinION using FLO-MIN106 Flow Cell, base-called reads with Albacore, and aligned the resulting reads to Human reference assembly hs37d5 using minimap2 -a -z 600,200 -x map-ont. BAM files were processed first with samtools calmd -e, then with a custom script, sam_to_event_matrix.py. The latter parsed the CIGAR strings and MD tags to collect rates of match, mismatch, insertion, and deletion. The samtools step was necessary to provide MD tags. While these alignments are likely to exclude (or underreport) TRs, which are frequently excluded from the reference genome (Sedlazeck *et al.*, 2018), we nonetheless expect the observed error profiles to be good representations of errors introduced by sequencing. We do not account for genuine polymorphism by which HG002 differs from the reference human genome, hence the profiles slightly overestimate error. However, with human average polymorphism levels of 0.1% (Sachidanandam *et al.*, 2001), this is unlikely to affect our estimates in any significant way.

#### Supplementary Note S3. Analysis of Simulated Reads and Tool Comparison

We tested NCRF on simulated data, comparing it to TRF (Benson, 1999) and Minimap2 (Li, 2018). A 3.3-Mb mock genome was created, which consisted of a mix of repeats and random sequence. Six primary repeat elements had periods of 5, 10, 20, 40, 80, and 171 bp. The period 5 element was $(AATGG)_n$, the 171-mer was alpha satellite, and the other four repeat elements were generated randomly (Table S4). A total of 2,000 repeat arrays were generated, with lengths drawn from a rounded normal distribution with a mean of 1,000 bp, a standard deviation of 400 bp, and a minimum length of 500 bp (lengths drawn from the distribution, but less than 500, were discarded). Half of the arrays were chosen from the six primary elements. The others are intended as false positive bait for processes searching for the primary elements. Bait arrays were formed by choosing, at random, a motif with edit distance 1 from one of the primary elements, and repeating this motif; each bait array was the result of another random choice. Bait arrays formed from higher edit distance motifs were not utilized, since distinguishing a motif from an edit distance 1 imposter is the worst case. Arrays were positioned randomly across the single-sequence "genome", with a minimum of 500 bp of random sequence between them.

Simulated PacBio and Nanopore reads were separately sampled from the mock genome. Reads were sampled from a minimum of 1 kb to a maximum of 20 kb, with probability density linearly decreasing across this range, for an expected average length of 7.3 kb, to crudely mimic the variation in read lengths seen in actual sequencing data. Sequencing depth was 5x. Sequencing errors were applied according to the derived profiles for the two technologies. The positions of the embedded repeats in both the genome and simulated reads were recorded, comprising the ground truth by which the tools were evaluated.

NCRF was run using technology-specific scoring derived from error profiles above (Table S3). Error rate limit was set to 25%, and repeats shorter than 500 bp were discarded. Consensus filtering (see Main Text and Supplementary Note S1C) was applied. The alignments were compared to the ground truth to calculate true positive (TP), false positive (FP), and false negative (FN) rates (see *Classifier Analysis*, below).

TRF was run (Supplementary Note S4) using scoring parameters shown in an example at the TRF command line use help page https://tandem.bu.edu/trf/trf.unix.help.html. The resulting alignments were parsed from the TRF's html output, and repeats shorter than 500 bp, or with a different repeat motif, were discarded.

For Minimap2, the simulated reads were mapped to an artificial reference genome consisting of a single array (for each motif) longer than the longest array in the mock genome, so that any noisy array has the potential to be aligned in its entirety Technology-specific parameterization (Supplementary Note S5) was used. Alignments with a mapping quality of zero were discarded. The remaining alignments for both TRF and Minimap2 were compared to the ground truth to calculate TP, FP, and FN rates.

##### Classifier analysis

Aligned results from each tool were compared to the known truth recorded when the simulated reads were generated. Each base reported by a tool was counted as either a TP, FP, overlapped true positive (TPO), or overlapped false positive (FPO). The latter two categories account for the fact that TRF and minimap will occasionally report overlapping intervals for the motif. When this occurred, a base occurring in more than one interval was counted as a TP or FP, and the remaining occurrences were counted as

TPO or FPO. The counts were used to compute the standard classifier statistics TPR and FDR. Results are shown in Table S2.

#### Supplementary Note S4. Tandem Repeats Finder

We augmented Tandem Repeats Finder (TRF) with custom post-processing to facilitate search for repeats of specific motifs. The first stage ran TRF on reads to find all candidate repeats. The second stage extracted the repeats of interest, converting them into a format suitable for classifier evaluation.

Our TRF command line was

```
trf reads.fa 2 7 7 80 10 100 205
```

Where

- 2 is matching weight
- 7 is mismatching penalty
- 7 is indel penalty
- 80 is match probability
- 10 is indel probability
- 100 is minimum score
- 205 is maximum period

Our tests indicated that a minimum score of 100 gave the best sensitivity for a length cutoff of 500 bp.

The maximum period of 205 was determined as 120% of the period of the longest search motif (171). This is necessary because TRF initially measures period as it is observed in the context of the sequenced reads. If a sequencer causes more insertions than deletions, the observed period will be longer than the period in the molecule. Increasing the maximum period parameter accommodates this effect.

We used TRF in html output mode. This was necessary to glean the base-by-base alignment details needed for classifier evaluation.

The second stage used a custom script, harvest_trf_html.py. In addition to parsing the html files, this script discards alignments with an unwanted consensus, and repeat arrays that are too short (aligned to less than 500 bp in the hypothetical molecule). The surviving alignments are output in a format similar to NCRF's alignment output.

We used Tandem Repeats Finder 4.09 for 64 bit Linux, downloaded Apr/30/2018 from https://tandem.bu.edu/trf/trf409.legacylinux64.download.html.

#### Supplementary Note S5. Minimap2

We asked Minimap2 to align reads to artificially created sequences specific to our repeats of interest. Custom post-processing then converted the output to a format suitable for classifier evaluation.

For a given motif, the artificial sequence consisted of a perfect copy of the motif 5% longer than the longest repeat array we wanted to find. For our simulation analysis, we knew the longest embedded arrays, which led to bait sequences of length 2,636 bp. Artificial sequences for all motifs were collected as separate sequences in a single fasta file.

Our Minimap2 command lines were

(for Pacbio) minimap2 -x map-pb --cs bait.fa reads.fa

(for ON) minimap2 -x map-ont --cs bait.fa reads.fa

The second stage used a custom script, minimap2_cs_to_events.py. This converts a paf file to a format similar to NCRF's alignment summaries. It also discards low-quality alignments. In our use we set --minquality=1, so that alignments with quality 0 (as defined in Minimap2's paf format) were discarded.

We used Minimap2 version 2.10-r761 for 64 bit Linux, downloaded May/30/2018 from https://github.com/lh3/minimap2/releases.

#### Supplementary Note S6. Analysis of Sequenced Human Reads

One application of NCRF is to find the length distribution of specific TRs in sequenced reads. To demonstrate the utility of NCRF for real sequencing data, we used it to find repeats of $(AATGG)_n$ in the human genome sequenced with PacBio and Nanopore technologies (Zook *et al.*, 2016; Jain *et al.*, 2018); data sources are provided below. We specifically searched for long repeats (at least 500 bp in length).

We observed different read length distributions in the two sequencing technologies, as well as different sequencing yield. Observed repeat lengths are affected by read lengths, primarily due to reads that partially cover a repeat (such repeats are then observed with a shorter length). In order to be able to meaningfully compare repeat lengths, we subsampled both sets of reads to achieve similar read length distributions.

First, the actual length distribution of all reads from each technology was computed. Counts for lengths were accumulated in bins of similar lengths (instead of exact lengths), since specific long lengths are rarely in both sets. Moreover, a logarithmic scale was used so that long reads had wider bins. The two binned distributions were compared to pick a maximal common distribution. Then, for each technology, reads were sampled according to this common distribution, with counts divided in rough proportion to abundance in the original read files. Figure S2 shows the resulting sampled distribution used, which resulted in a total of 16.5 Gb of the sequenced data for each technology.

The NCRF software distribution contains an example script, common_length_distribution.sh, to demonstrate the process.

#### Supplementary Note S7. Preliminary Consideration of Other Tools

Before embarking on the evaluations presented in Supplementary Note S3, we performed brief experiments with several other tools to determine whether it was feasible to include them.

It should be noted that none of the tools were suitable for easy comparison right off the shelf. Each required some additional data manipulation for input or, more commonly, for output. The decision to include a tool was partially motivated by the amount of programming effort necessary to extract, from the tool's output, the information relevant to the evaluation.

##### 7A. HMMER

We tried HMMER (Eddy, 1998) to search for motifs of length 5, 10, 20, 40, 80, and 171 bp. For each motif, 10 reads were chosen from simulated reads sampled from a 3.3-Mb mock genome, reads known to include segments covering a repeat of the motif in the mock genome. Supplementary Note S3 provides details on the creation of the genome and sampling of reads.

For each motif, hmmbuild was used to build an hmm file from an "msa" file containing the motif and its reverse complement. Hmmsearch was then used to find matches for the hmm file in the 10 reads. For simulated reads from both technologies, it performed well on the 40-mer and longer, reporting at least 80% of the motif copies known to be present. However, it found no copies of the 5-mer, (over 2,000 copies were present in each technology), and it found less than 2%, of the 10-mers (over 1,000 present).

After trying a few other preliminary tests, we concluded that HMMER was not suitable for finding short motifs, and did not test it further.

We used HMMER version 3.1b2, downloaded Oct/5/2017 from http://hmmer.org.

##### 7B. Tantan

Tantan (Frith, 2011) is a soft masking tool that marks novel repeated motifs in genomic sequence. Its design was motivated by a need to exclude repeats during alignment of sequenced reads to an assembled genome.

In early tests, searching for $(AATGG)_n$ in both simulated and real data, Tantan identified significantly less repeated content compared to NcRF. This was improved by parameter tweaking — for example, by enabling indels (which are disabled by Tantan's defaults). But even after this effort, Tantan identified less than half of the repeated content compared to NcRF.

It was observed that Tantan does not allow for penalizing insertions differently than deletions. Moreover, when a repeat array is identified it is only annotated by marking its sequence in lowercase. It doesn't report the putative motif, nor does it provide an alignment of the interval to itself from which a consensus motif could be easily derived. While this information could have been derived by post-processing using another aligner, doing so would have raised the question of whether the aligner used was the best for this purpose.

Thus it would have been difficult to perform the classifier analysis (Supplementary Note S3) without significant additional work, and given the poor comparative results of the test we did do, we decided not to include Tantan in out evaluation.

**7C. PacmonSTR**

PacmonSTR (Ummat and Bashir, 2014) is designed to identify TRs and estimate the number of TR elements in long sequenced reads. However, it requires an annotated reference genome indicating the TRs of interest.

Input to PacmonSTR is alignments between reads and the TRs in the reference genome. Alignments are generated externally, but realigned by an internal HMM. Alignments must include flanking regions — the flanking regions are used to infer an error model for the HMM that realigns the TR.

This adaptive realignment, adapting to the characteristics of the locality of the discovered element, is a great feature. However, the requirement of knowing the flanks of a TR make it unsuitable for discovering repeats to occur in unassembled segments of the genome.

**7D. Alpha-CENTAURI**

Alpha-CENTAURI (Sevim et al. 2016) is designed to identify TRs in long sequenced reads, and specifically to identify higher order repeat structures.  Since this tool relies on HMMER (Eddy, 1998) to find locations of the motif (without regard to repetition), we deferred evaluation it to our evaluation of HMMER (Supplementary Note 7A).

**7E. Replong**

While designed to find *repeats* in long sequenced reads, RepLong (Guo *et al.*, 2018) doesn't search for tandem repeats, but rather looks for repeat *families*. Segments of reads are considered similar if they have similar length and similar content, regardless of whether or not the segments contains a TR. Thus a 2 kbp segment consisting solely of a repeated 5-mer TR is identified as similar only to other segments that contain the same TR *and* with length close to 2 kbp.

**Data sources:**

[GIAB-PacBio-reads] HG002 PacBio reads. Files are linked from the table at this URL. Only the files with NIST_SAMPLE_NAME=HG002 are data for HG002.
https://github.com/genome-in-a-bottle/giab_data_indexes/blob/master/AshkenazimTrio/sequence.index.AJtrio_PacBio_MtSinai_NIST_hdf5_10102018

[GIAB-Nanopore-reads] HG002 Oxford Nanopore reads.
ftp://ftp-trace.ncbi.nlm.nih.gov/giab/ftp/data/AshkenazimTrio/HG002_NA24385_son/Ultralong_OxfordNanopore/combined_2018-05-18/combined_2018-05-18.fastq.gz

#### SUPPLEMENTARY FIGURES

**Figure S1.** Typical workflow. Noise-Cancelling Repeat Finder (NCRF) is given a set of repeat motifs and sequence files, and reports alignments. Alignment files are post-processed by scripts included in the package and then converted to a tabular text format to undergo further processing (e.g., filtering for overlaps, see Supplementary Note S1E, or user-developed scripts).

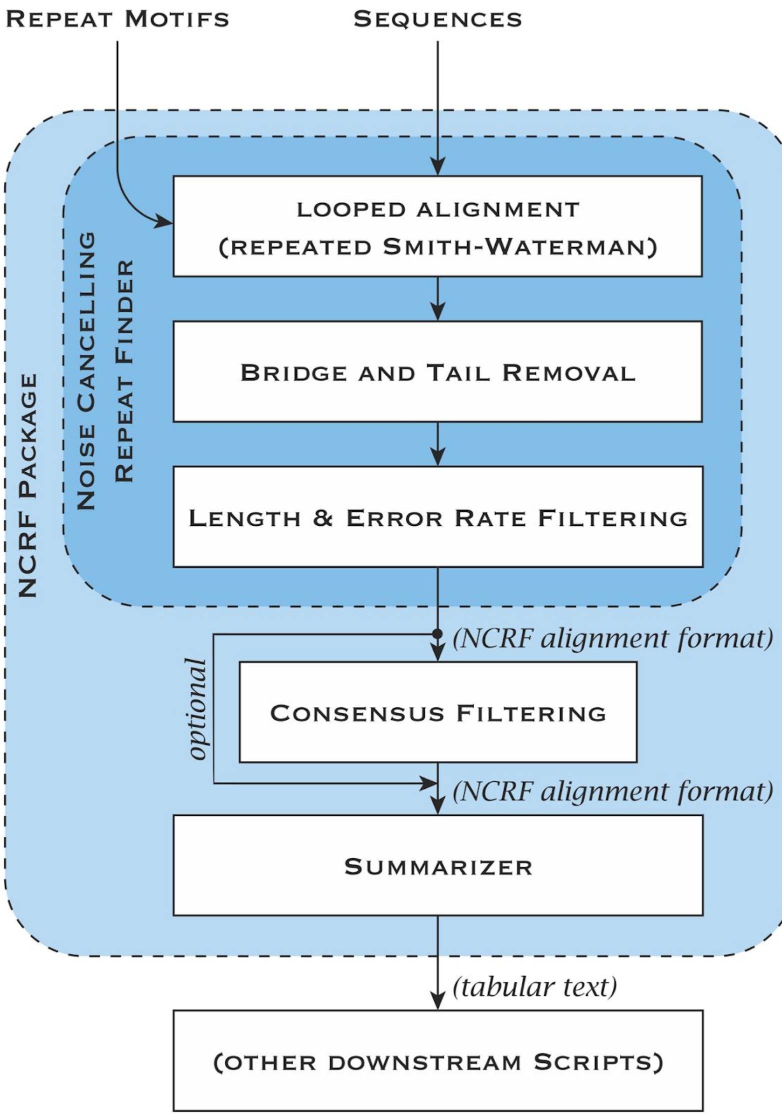

**Figure S2.** Distribution of read lengths (in base pairs) in PacBio and Nanopore reads. Both were subsampled to achieve a similar distribution with 16 Gb (mean read length of 10.6 kb).

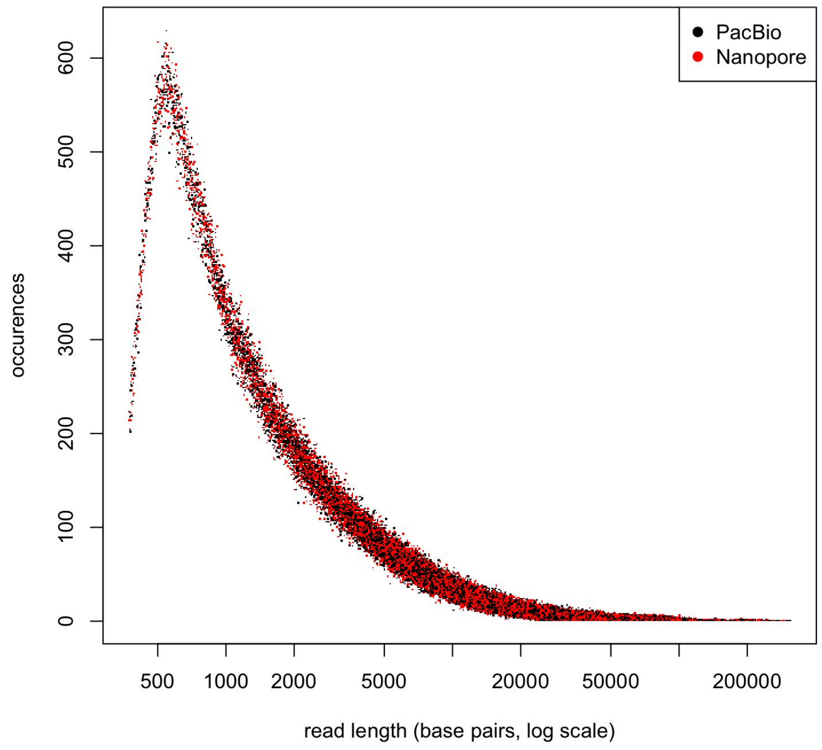

#### SUPPLEMENTARY TABLES

**Table S1. Observed error rates in PacBio and Nanopore alignments.** m=matches, mm=mismatches, io=insertion open, ix=insertion extend, do=deletion open, dx=deletion extend. Overall error rates are measured as 1 - m/(m+mm+io+ix+do+dx).

| Technology | Error Rate | m | mm | io | ix | do | dx |
|---|---|---|---|---|---|---|---|
| PacBio | 14.90% | 203521495440 | 4039815573 | 15516830204 | 5868396512 | 8622457071 | 1709970892 |
| | | 85.06% | 1.69% | 6.48% | 2.45% | 3.60% | 0.71% |
| ON | 16.10% | 17603088986 | 965110092 | 492506288 | 299904008 | 702350550 | 919627709 |
| | | 83.89% | 4.60% | 2.35% | 1.43% | 3.35% | 4.38% |

**Table S2. Classifier analysis results.** TP=true positives, FP=false positives, TPO=additional overlapped true positives, FPO=additional overlapped false positives, TPR=true positive rate, FDR=false discovery rate.

##### A. PacBio

| Tool | Motif | TP | FP | TPO | FPO | TPR | FDR |
|---|---|---|---|---|---|---|---|
| NCRF unfiltered | AATGG | 927273 | 27542 | 0 | 0 | 99.02% | 2.89% |
| NCRF with consensus filtering | AATGG | 931312 | 6903 | 0 | 0 | 98.59% | 0.74% |
| TRF | AATGG | 926817 | 6289 | 0 | 0 | 98.97% | 0.67% |
| minimap2 | AATGG | 727132 | 258087 | 8306 | 3840 | 77.65% | 26.20% |
| NCRF unfiltered | motif10 | 883248 | 572951 | 0 | 0 | 99.00% | 39.35% |
| NCRF with consensus filtering | motif10 | 884284 | 5829 | 0 | 0 | 98.07% | 0.66% |
| TRF | motif10 | 881909 | 6343 | 1136 | 2 | 98.85% | 0.71% |
| minimap2 | motif10 | 641655 | 98624 | 248 | 0 | 71.92% | 13.32% |
| NCRF unfiltered | motif20 | 807807 | 782079 | 0 | 0 | 99.29% | 49.19% |
| NCRF with consensus filtering | motif20 | 807807 | 9909 | 0 | 0 | 98.29% | 1.21% |
| TRF | motif20 | 793318 | 6335 | 0 | 0 | 97.51% | 0.79% |
| minimap2 | motif20 | 589521 | 182864 | 3532 | 0 | 72.46% | 23.68% |
| NCRF unfiltered | motif40 | 850628 | 1020783 | 0 | 0 | 99.43% | 54.55% |
| NCRF with consensus filtering | motif40 | 848600 | 14216 | 0 | 0 | 98.16% | 1.65% |
| TRF | motif40 | 754358 | 4356 | 0 | 0 | 88.18% | 0.57% |
| minimap2 | motif40 | 553426 | 577921 | 1221 | 1224 | 64.69% | 51.08% |
| NCRF unfiltered | motif80 | 854737 | 864383 | 0 | 0 | 99.20% | 50.28% |
| NCRF with consensus filtering | motif80 | 689336 | 14266 | 0 | 0 | 78.67% | 2.03% |
| TRF | motif80 | 564275 | 5119 | 0 | 0 | 65.49% | 0.90% |
| minimap2 | motif80 | 480047 | 612957 | 2868 | 3250 | 55.71% | 56.08% |
| NCRF unfiltered | alpha-satellite | 863704 | 862798 | 0 | 0 | 99.44% | 49.97% |
| NCRF with consensus filtering | alpha-satellite | 138836 | 5068 | 0 | 0 | 15.76% | 3.52% |
| TRF | alpha-satellite | 43669 | 27 | 0 | 0 | 5.03% | 0.06% |
| minimap2 | alpha-satellite | 494941 | 569334 | 1041 | 3816 | 56.98% | 53.50% |

#### B. Nanopore

| Tool | Motif | TP | FP | TPO | FPO | TPR | FDR |
|---|---|---|---|---|---|---|---|
| NCRF unfiltered | AATGG | 875822 | 3656 | 0 | 0 | 95.37% | 0.42% |
| NCRF with consensus filtering | AATGG | 875822 | 3656 | 0 | 0 | 94.61% | 0.42% |
| TRF | AATGG | 906248 | 3989 | 0 | 0 | 98.68% | 0.44% |
| minimap2 | AATGG | 699551 | 19759 | 719 | 0 | 76.17% | 2.75% |
| NCRF unfiltered | motif10 | 843491 | 42037 | 0 | 0 | 94.88% | 4.75% |
| NCRF with consensus filtering | motif10 | 843970 | 3848 | 0 | 0 | 94.03% | 0.45% |
| TRF | motif10 | 866352 | 5407 | 2043 | 0 | 97.45% | 0.62% |
| minimap2 | motif10 | 705864 | 40481 | 2445 | 0 | 79.40% | 5.42% |
| NCRF unfiltered | motif20 | 795937 | 374867 | 0 | 0 | 94.83% | 32.02% |
| NCRF with consensus filtering | motif20 | 795937 | 8158 | 0 | 0 | 93.76% | 1.02% |
| TRF | motif20 | 764439 | 7052 | 0 | 0 | 91.08% | 0.91% |
| minimap2 | motif20 | 580872 | 486081 | 1328 | 2641 | 69.21% | 45.56% |
| NCRF unfiltered | motif40 | 819919 | 853463 | 0 | 0 | 95.18% | 51.00% |
| NCRF with consensus filtering | motif40 | 797986 | 29730 | 0 | 0 | 91.85% | 3.59% |
| TRF | motif40 | 393475 | 3396 | 0 | 0 | 45.68% | 0.86% |
| minimap2 | motif40 | 477715 | 742383 | 455 | 3373 | 55.46% | 60.85% |
| NCRF unfiltered | motif80 | 880476 | 769318 | 0 | 0 | 95.90% | 46.63% |
| NCRF with consensus filtering | motif80 | 604307 | 13485 | 0 | 0 | 65.27% | 2.18% |
| TRF | motif80 | 133740 | 61 | 0 | 0 | 14.57% | 0.05% |
| minimap2 | motif80 | 578553 | 663602 | 2665 | 6798 | 63.01% | 53.42% |
| NCRF unfiltered | alpha-satellite | 791017 | 869031 | 0 | 0 | 94.96% | 52.35% |
| NCRF with consensus filtering | alpha-satellite | 50844 | 1304 | 0 | 0 | 6.02% | 2.50% |
| TRF | alpha-satellite | 0 | 0 | 0 | 0 | 0.00% | — |
| minimap2 | alpha-satellite | 509384 | 624405 | 1220 | 4181 | 61.15% | 55.07% |

**Table S3. NCRF alignment scoring parameters.** Default scoring parameters implemented in NCRF, as derived from observed error models. m=match reward, mm=mismatch penalty, io=insertion open penalty, ix=insertion extend penalty, do=deletion open penalty, dx=deletion extend penalty.

| Technology | m | mm | io | ix | do | dx |
|---|---|---|---|---|---|---|
| PacBio | 10 | 35 | 33 | 21 | 6 | 28 |
| Nanopore | 10 | 63 | 51 | 98 | 27 | 34 |

**Table S4. Primary repeat elements in the mock genome.** The alpha satellite sequence originates from (Sevim *et al.*, 2016).

| | |
|---|---|
| AATGG | AATGG |
| motif10 | CACTGCTGGT |
| motif20 | GTTAGCGGTCGTGCTGATGG |
| motif40 | GGCTCCTATCTCGCCTGTTCCCGGGTTCCTCTTATTCTCA |
| motif80 | GAGATTGGAGTCCAAGAAATTCAGTCACCTTTCAGCGGTTCCAGTCACGGCGCTAAGTG CCTATTGACCCGCTACTGTTT |
| alpha | TCTGTCTAGTTTTTATATGAAGATATTCCCTTTTCCACCATAATCCTCAAAGCGCTCCAAAT ATCCACTTGCAGATTCTACAAAAAGAGTGTTTCCAAACTGCTCTATCAAAAGAAATGTTCA ACTCTGTGAGTTGAATACACACATCACAAAGAAGTTTCTGAGAATGCT |
